# An allosteric ligand stabilizes distinct conformations in the M2 muscarinic acetylcholine receptor

**DOI:** 10.1101/2021.02.14.431178

**Authors:** Jun Xu, Harald Hübner, Yunfei Hu, Xiaogang Niu, Peter Gmeiner, Changwen Jin, Brian Kobilka

## Abstract

Allosteric modulators provide therapeutic advantages over orthosteric drugs. A plethora of allosteric modulators have been identified for several GPCRs, particularly for muscarinic receptors (mAChRs)^1,2^. To study the molecular mechanisms governing allosteric modulation, we utilized a recently developed NMR system to investigate the conformational changes in the M2 muscarinic receptor (M2R) in response to the positive allosteric modulator (PAM) LY2119620. Our studies provide the first biophysical data showing that LY2119620 can substantially change the structure and dynamics of M2R in both the extracellular and G-protein coupling domains during the activation process. These NMR data suggest that LY2119620 may function by stabilizing distinct sets of conformations not observed in the presence of orthosteric agonists alone, which may account for the different signaling behaviors of the M2R when bound to LY2119620. Our studies provide new structural information for understanding the mechanism of GPCR allostery, and may facilitate the rational design of allosteric therapeutics targeting muscarinic receptors.

## Introduction

Allosteric modulators of GPCRs are defined as small molecules that can bind to an allosteric site that is topographically distinct from the endogenous ligand binding site (also known as the orthosteric site)^1,3,4^. Recent X-ray crystallographic studies have characterized several distinct allosteric binding sites in GPCRs^1,5–7^. The mAChRs, and in particular the M2 subtype, are among the best-characterized Family A GPCRs for understanding the allosteric modulation by small molecules^2,8,9^. In addition to the wide spectrum of orthosteric modulators, many allosteric modulators with varying pharmacological profiles have been characterized for all mAChR subtypes in the past decades^4,8,10–12^. These allosteric modulators can regulate the affinity and/or efficacy of orthosteric ligands, and some of them can modulate the receptor activity on their own; therefore, they may provide us a means to fine-tune the biological responses to the natural signaling patterns and help us to achieve the functional selectivity or signaling bias^3,13,14^. In addition, because the allosteric site shows greater sequence divergence among receptor subtypes in comparison to residues comprising the orthosteric pocket^15^, a better understanding of mAChRs allostery could facilitate the discovery of subtype selective ligands.

Previous studies have characterized the extracellular vestibule (ECV) as one of the most common allosteric binding sites for mAChRs ^2,16,17^. The recent structures of active M2R in complex with the high efficacy orthosteric agonist, iperoxo (Ixo), in the presence or absence of the positive allosteric modulator (PAM), LY2119620, represented a major advance in understanding allostery for a mAChR at an atomic level^7^. However, the static structures only represent the first step in our understanding of mechanisms underlying GPCR allostery as they cannot describe the dynamic characteristics of the receptor. Moreover, both structures were stabilized in the active-state by a nanobody (Nb9-8) that binds to the cytoplasmic surface of the M2R, making it difficult to observe structural differences at the cytoplasmic end of TM segments due to binding of the PAM. Recent atomic molecular dynamics simulations studies have contributed to our understanding of the mechanisms of the cooperativity between a series conventional negative allosteric modulators (NAMs) and orthosteric antagonists^17^, whereas less is known about how PAMs modulate structure and dynamics of M2R bound to orthosteric agonist.

Using ^13^CH_3_-ε-methionine (^13^CH_3_-ε-M) NMR, we recently investigated the conformational dynamics of M2R bound to distinct orthosteric modulators, and provided the first biophysical data demonstrating the highly dynamic nature of the M2R^18^. Here, we extend these studies to LY2119620, which is a recently identified PAM for the M2 muscarinic receptor^7,19^. In this study, we focus on characterizing the cooperative effects between LY2119620 and two orthosteric agonists, acetylcholine (ACh) and iperoxo (Ixo), on the M2R by monitoring the chemical environment changes surrounding five ^13^CH_3_-ε-M distributed throughout TM core of the receptor (Fig. S1)^18^.

## Results

### Pharmacology of LY2119620

Previous functional studies show that LY2119620 displays modest allosteric agonism on its own and can enhance agonist potency (affinity) when measured by [^35^S]GTPγS-binding and ERK1/2 phosphorylation assays^7,19^. In the present study, we performed G-protein IP-one accumulation and β-arrestin recruitment assays to evaluate the effects of LY2119620 on signaling behaviors of the neurotransmitter acetylcholine (ACh) and and the agonist iperoxo (Ixo) under the same conditions (Fig. 1, Table S1). Our results show that, in contrast with previous data^19^, LY2119620 displays a partial agonist activity by itself in both G-protein and β-arrestin pathways. We observed much higher efficacy of LY2119620 alone (60% of maximal ACh response) in these two assays than previously measured using [^35^S]GTPγS binding (20% of ACh response)^19^. This is probably due to the greater sensitivity of the G-protein IP-one and β-arrestin recruitment assays. As shown by the IP-one accumulation data, we found that the highest concentrations of LY2119620 could inhibit the maximal G-protein activation in response to both orthosteric agonists, most notable for Ixo (Fig.1**a, b, e, f**). In contrast to the previous data measured using ERK1/2 phosphorylation assay, we observed significant enhancement of the efficacy (Emax) by 20-30% in the β-arrestin recruitment, especially for ACh (Fig.1**c, d, g, h**). The discrepancy with previous studies^7,19^ is likely due to the fact that ERK1/2 phosphorylation involves both G-protein and β-arrestin signaling pathways^20^.

**Fig. 1.**
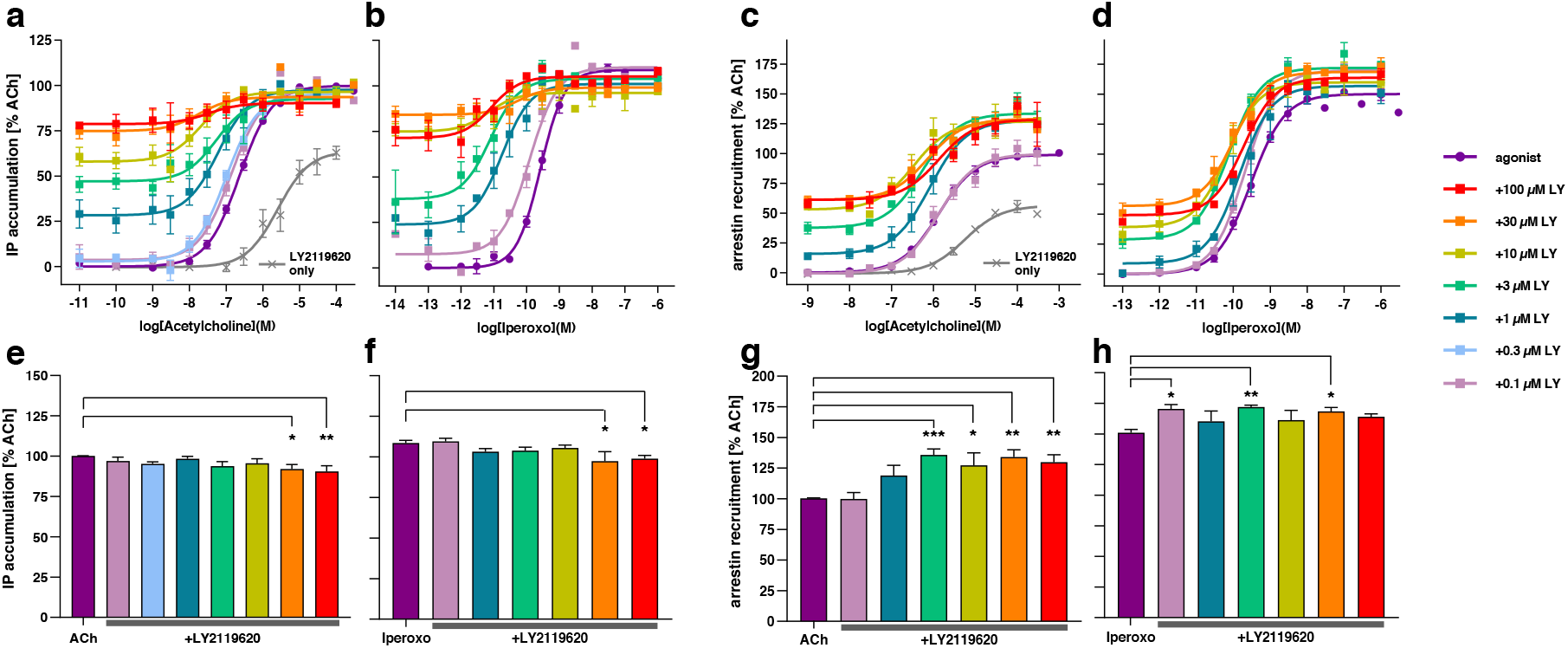
Pharmacology of LY2119620. (a-d) Concentration-response curves of acetylcholine (a and c) and iperoxo (b and d) toward G protein activation and β-arrestin-2 recruitment in the presence of different concentrations of LY2119620. (e-h) Statistics of Emax calculations for G protein activation and β-arrestin-2 recruitment. G protein activation and β-arrestin-2 recruitment are measured by a cell-based Gqi-inositol phosphate accumulation assay and an enzyme fragment complementation based arrestin recruitment assay, respectively. Data are presented as mean ± SEM of three to fourteen independent experiments with repeats in duplicate. Emax values were analyzed by One-way ANOVA applying Dunnett’s multiple comparisons in PRISM 8.0. The significance (p < 0.05) is displayed for the efficacy of a distinct concentration of LY2119620 versus the agonist alone. Mean values were calculated as Emax ± SEM [% relative to ACh].

### Effects of LY2119620 on the HSQC spectrum of apo-state M2R

As noted above, we previously used NMR spectroscopy to study conformational changes and dynamics around five ^13^CH_3_-ε-M distributed throughout the transmembrane bundle (Fig. S1**a**). Crystal structures show that these ^13^CH_3_-ε-M are well positioned to detect the local conformational changes. As shown in Fig. S1, M406^6.54^ is located at the top of TM6, where large structural changes occurre upon receptor activation and this position is close to the allosteric pocket. Thus M406^6.54^ is sensitive to conformational changes involving allosteric ligand binding (Fig. S1**b**). M77^2.58^ is located at the extracellular half of TM2, the close proximity (within 5Å) to aromatic residues Y430^7.43^ and Y80^2.61^ that are part of the orthosteric pocket and allosteric site (Fig. S1**c**) makes M77^2.58^ sensitive to changes in these regions. M112^3.41^ is found just below the ligand binding pocket in TM3 and close to the conserved P^5.50^-I/V^3.40^-F^6.44^ motif, a conformational switch in several Family A GPCRs, thus M112^3.41^ is positioned to detect the propagation of conformational changes from the ligand binding pocket to the cytoplasmic surface (Fig. S1**d**). M143^4.45^ is located at the cytoplasmic side of TM4 (Fig. S1**d**), and it is close to the TM3 cytoplasmic domain and ICL2, which have been found to interact with with G proteins^21–23^. Of interest, the cavity formed by TM3/ICL2/TM4 was recently found to be a binding site of certain allosteric modulators^24^. M202^5.54^ is located between the P^5.50^-I/V^3.40^-F^6.44^ motif and cytoplasmic end of TM5, with the ε-^13^CH^3^ group positioned between TM5 and TM6 (Fig. S1**e**). M202^5.54^ is therefore an excellent sensor to monitor conformational changes in TM5 and TM6.

The overall spectra of the M2R bound to different ligands or ligand combinations are summarized in Fig. S2. Crystal structures reveal that LY2119620 binds to the contracted extracellular vestibule, which is stabilized by agonists (Fig. 2**a**). We find that LY2119620 alone stabilizes a different conformation relative to the apo-state receptor (Fig. 2**b**), indicating that even without agonist, LY2119620 is able to bind to the receptor. This is in agreement with it’s partial agonist activity (Fig. 1).

**Fig. 2.**
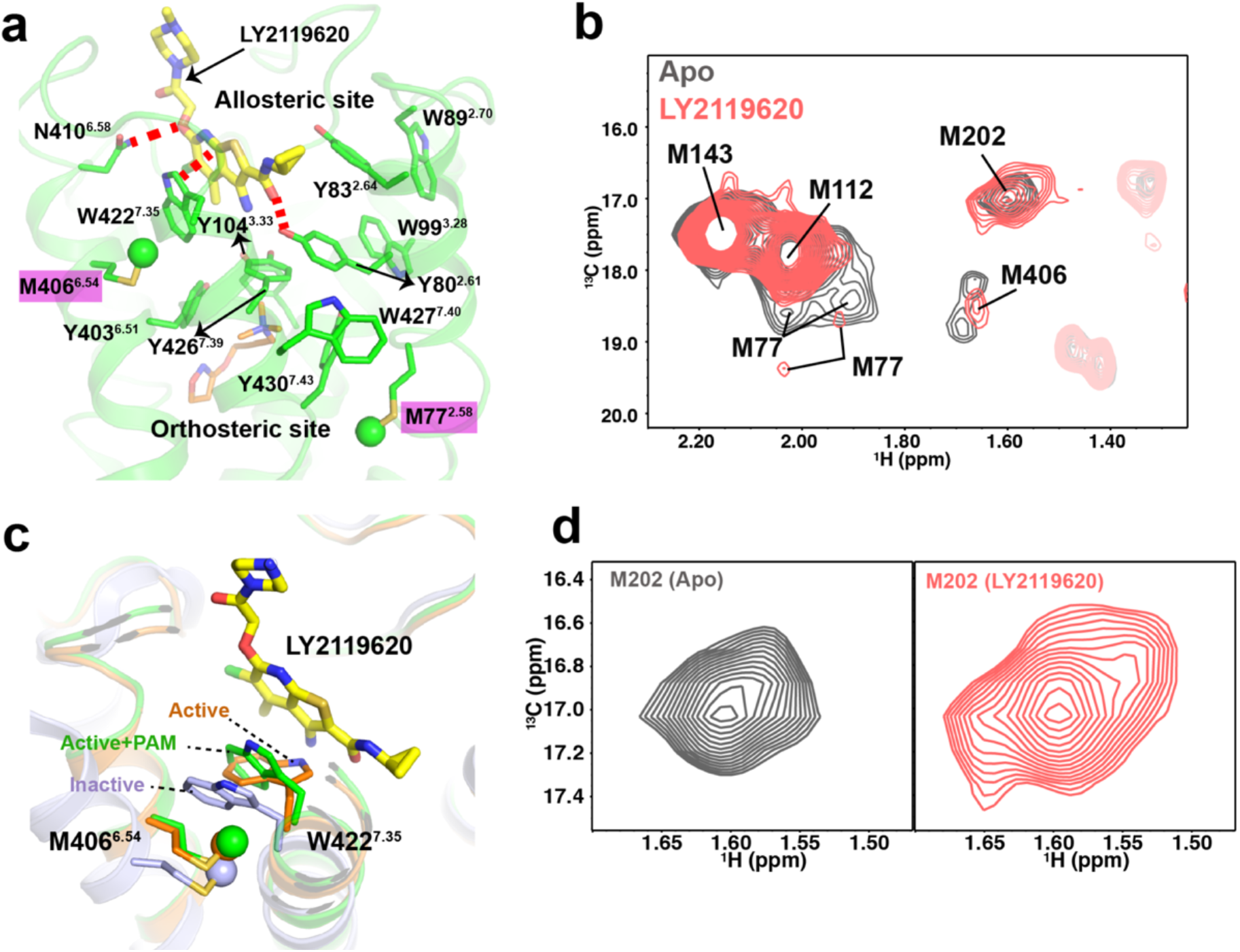
Effects of LY2119620 on the structure and dynamics of apo-state M2R. (a) Interaction between LY2119620 and the M2R extracellular vestibule (PDB: 4MQT). Polar contacts and aromatic stacking interactions are highlighted with red dashed lines. The residues constituting the aromatic network between M77^2.58^ and M406^6.54^ are shown in sticks. (b) Comparison of overall HSQC spectra between apo-state M2R and LY2119620-bound M2R. (c) Superimposition of W422^7.35^. sidechain conformations in inactive (light blue, 3UON), active (orange, 4MQS) and active/PAM (green, 4MQT) structures. M406^6.54^ located close to W422^7.35^ is shown in sticks. (d) Enlarged spectra of M202^5.54^ in apo-state and LY2119620-bound state.

The largest difference upon LY2119620 binding is in the peaks representing M77^2.58^, which is located adjacent to the orthosteric binding pocket (Fig. 2**a**). In the apo receptor, M77^2.58^ is represented by two peaks. Upon binding LY2119620 we still observe two weak peaks for M77^2.58^ one of which shifts significantly to a downfield position (Fig. 2**b**), suggesting a change of conformational dynamics of TM2, probably due to the slow to intermediate conformational exchange between at least two new sub-states. Crystal structures show that LY2119620 has direct interaction with Y80^2.61^ through a hydrogen bond in the presence of Ixo, while the interaction may be too weak to stabilize TM2 into a uniform conformation when no orthosteric agonist is bound (Fig. 2**a**). Since M77^2.58^ is positioned just below Y80^2.81^ and in close proximity to an aromatic network involving Y80^2.81^, W427^7.40^ and Y430^7.43^ (Fig 2**a**), the spectral changes of M77^2.58^ thus may reflect the dynamic character of Y80^2.61^. We previously found that M406^6.54^ would shift significantly to a downfield position in ^1^H dimension upon agonist binding (Fig. S2**a-c**)^18^, which may represent the formation of closed conformation (active) of the extracellular vestibule (ECV). However, when bound to LY2119620 alone, the M406^6.54^ peak shares the similar position in ^1^H dimension as the apo-state or inactive state. It’s hard to judge whether the ECV is open or closed under this condition, because the rotamer of W422^7.35^ in the LY2119620 bound active structure is similar to the rotamer in the inactive-state structure (Fig. 2**c**). The small chemical shift change of M406^6.54^ may be mainly due to the ring current effect from W422^7.35^ alone, instead of the whole extracellular region. The observation of a single weak peak for M406^6.54^ in LY2119620 bound relative to the two asymmetric peaks of apo-state in ^13^C dimension suggests different dynamics of the extracellular side of TM6.

The peaks for M112^3.41^ and M143^4.45^ are not affected by LY2119620 binding, suggesting little effect on the overall conformation and dynamics of TM3 and TM4 (Fig. 2**b**). This is probably due to the fact that LY2119620 does not have direct interaction with TM3 and TM4 (Fig. 2**a**). In contrast, we find that the peak corresponding to M202^5.54^ appears to be more heterogeneous (seems to become 3 peaks) than the apo-state peak (Fig. 2 **b** and **d**), indicating that LY2119620 binding might be able to alter the dynamic behavior of TM5-TM6 interface on the intracellular side (Fig. S1**e**), possibly by slowing down the chemical exchange between different states or by stabilizing different conformations. Although LY2119620 has no direct interaction with TM5, it directly interacts with TM6 and TM7 through hydrogen bond and aromatic stacking (Fig. 2**a**), which probably affects dynamic behavior of TM6 and TM7 on the intracellular side, and then propagates to the ^13^CH_3_-ε-M of M202^5.54^, which is positioned at the interface of TM5 and TM6 (Fig. S1**e**). As conformational changes involving TM5, 6 and 7 are required for receptor activation, these spectral changes may reflect structural changes responsible for the allosteric partial agonism of LY2119620.

### Effects of LY2119620 on the HSQC spectra of agonist-bound M2R

In addition to its direct allosteric agonism, LY2119620 can also affect the signalling behaviors of ACh and Ixo (Fig. 1**a-d**). We therefore obtained the HSQC spectra of LY2119620 and agonist-bound receptor. Interestingly, LY2119620 appears to significantly change the spectra of agonist-bound M2R in a quite complicated manner, rather than simply shifting the agonist-bound receptor to a more active state (Fig. 3). In the extracellular domain, the peaks corresponding to M77^2.58^ and M406^6.54^ for M2R bound to both LY2119620 and agonist are more similar to those observed for M2R bound to LY2119620 alone than M2R bound to agonist alone (Fig. 3**a-d**). As discussed above, the peak position for M406^6.54^ is likely due to the orientation of W422^7.35^ relative to M406^6.54^. In the inactive and iperoxo + LY2119620-bound crystal structures, M406^6.54^ faces the plane of W422^7.35^, while in the iperoxo bound structure, M406^6.54^ faces the edge of W422^7.35^ (Fig. 2**c**). The peaks for both M77^2.58^ and M406^6.54^ residues become more heterogeneous or weaker when compared with M2R bound to agonist alone, suggesting different conformational dynamics in the extracellular domain when bound to agonist and LY2119620. Taken together, these data suggest that LY2119620 and the orthosteric agonist can act cooperatively to stabilize distinct conformations and dynamics of the extracellular domain. The fact that LY2119620 plus agonist does not stabilize the same state as agonist alone may explain the fact that high concentrations LY2119620 suppress the maximal G-protein activation and enhances β-arrestin recruitment for both ACh and iperoxo (Fig. 1).

**Fig. 3.**
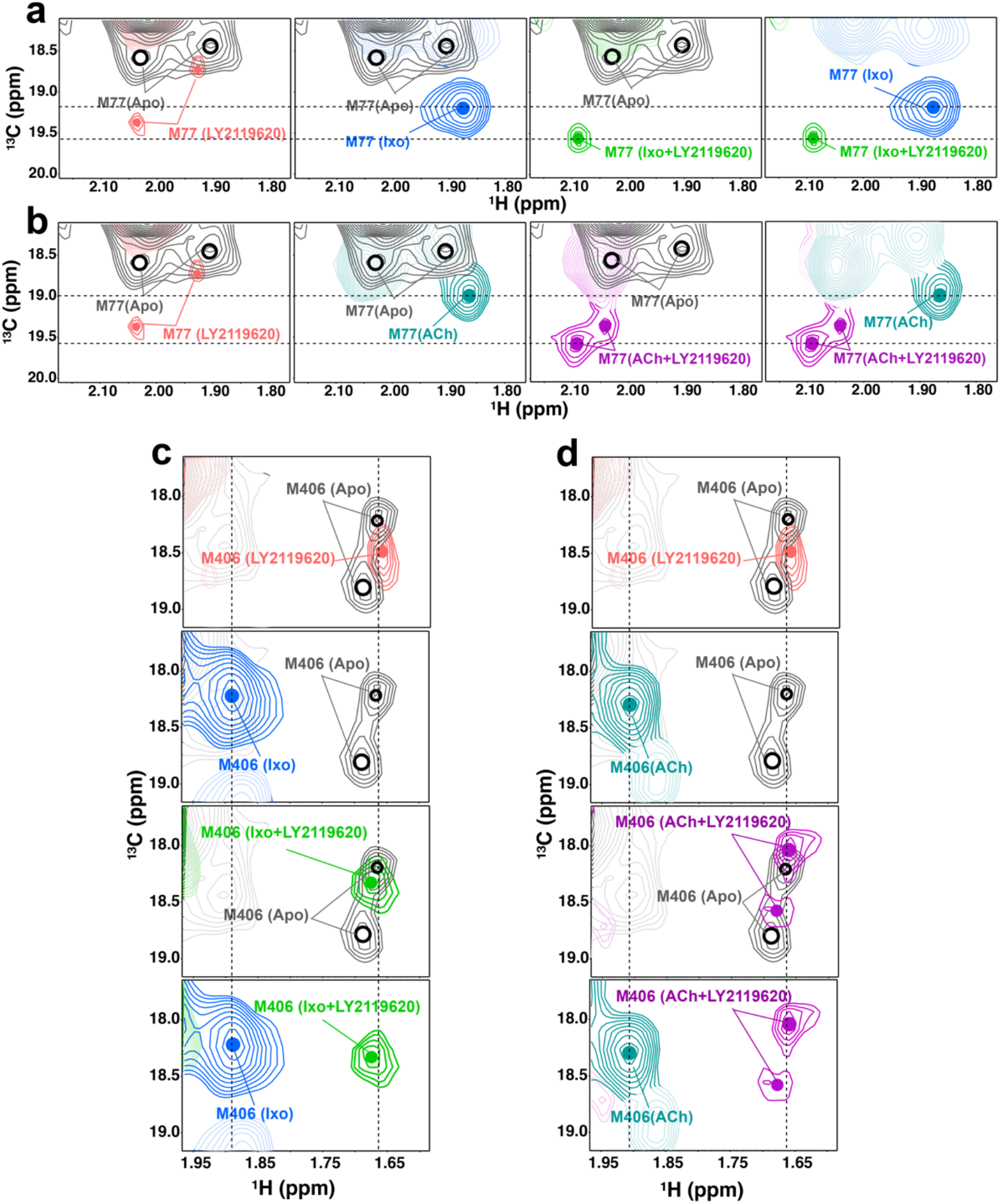
Effects of LY2119620 on the ^1^H-^13^C HSQC spectra of M77^2.58^ and M406^6.54^. The HSQC spectra of M77^2.58^ (a and b) and M406^6.54^ (c and d) in agonist (Ixo or ACh)-bound or agonist (Ixo or ACh) + LY2119620 bound state in comparison with the apo-state or LY2119620-bound state.

The effects of LY2119620 on ACh-bound M112^3.41^ and M143^4.45^ peaks are relatively small (Fig. 4). In contrast, LY2119620 has more significant effects on the spectra of Ixo-bound M112^3.41^ and M143^4.45^, and both spectra seem to split into two or multiple peaks, indicating the appearance of slow to intermediate conformational exchange for TM3 and TM4 (Fig. 4). In the presence of LY2119620, the Ixo-bound receptor the peaks representing M112^3.41^ and M143^4.45^ become more similar to ACh-bound receptor. These results indicate that LY2119620 may preferentially bind to the ACh-bound conformation over the Ixo-bound conformation in the absence of the intracellular signaling partner.

**Fig. 4.**
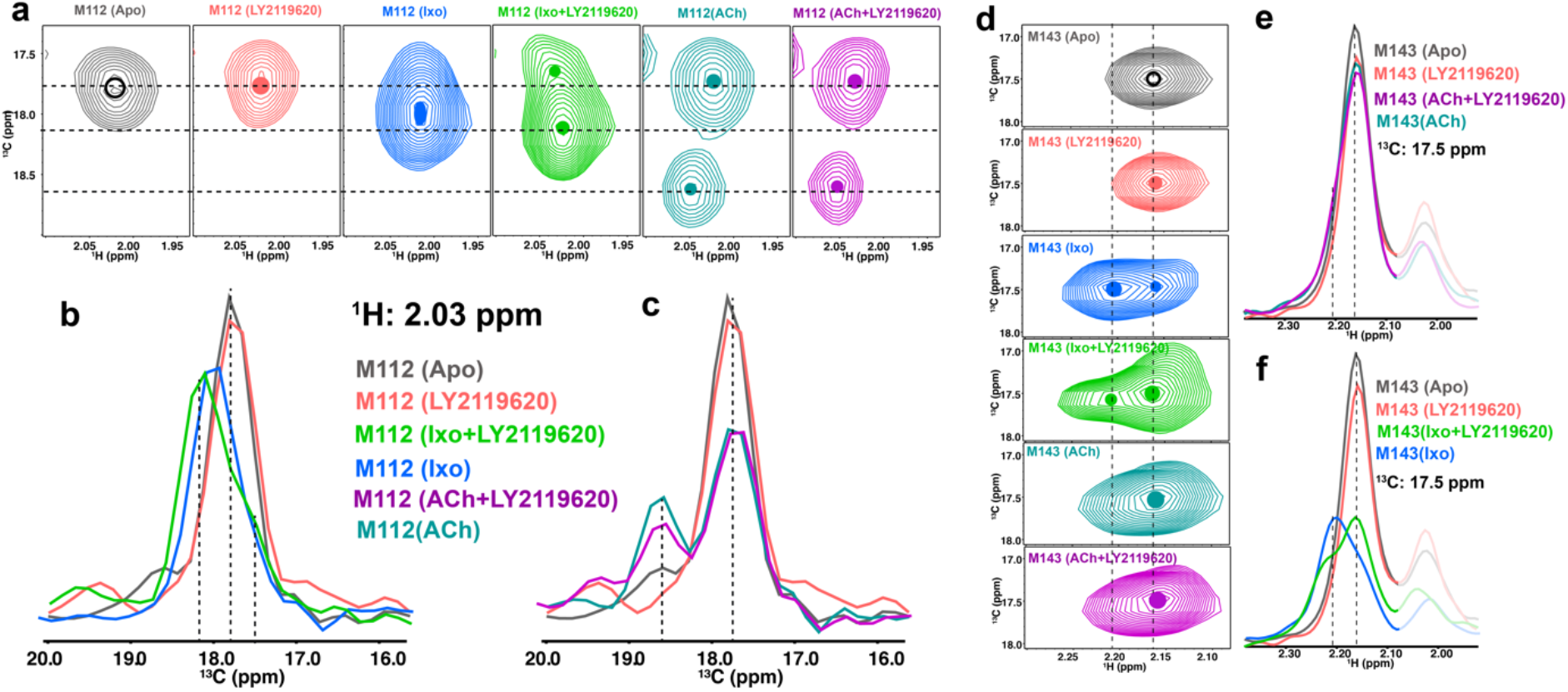
Effects of LY2119620 on the HSQC spectra of M112^3.41^ and M143^4.45^. (a) The 2D spectra of M112^3.41^ in different conditions: apo-state, LY2119620-bound state, iperoxo-bound state, iperoxo plus LY2119620-bound state, ACh-bound state and ACh plus LY2119620-bound state. (b and c) 1D slice analysis for the M112^3.41^ spectra in panel. 1D slices are extracted at 2.03 ppm in ^1^H dimension. (d) The 2D spectra of M143^4.45^ in different conditions: apo-state, LY2119620-bound state, iperoxo-bound state, iperoxo plus LY2119620-bound state, ACh-bound state and ACh plus LY2119620-bound state. (e and f) 1D slice analysis for the M143^4.45^ spectra in panel. 1D slices are extracted at 17.5 ppm in ^13^C dimension.

The effects of LY2119620 on the spectra of the intracellular residue M202^5.54^ in agonist-bound M2R are more pronounced than the effects of LY2119620 on the apo-state receptor (Fig. 2**d** and Fig. 5**a-b**). As discussed above, conformational changes may be propagated from the direct interaction of LY2119620 with TM6 and TM7. As we can see in Fig. 5**a-b**, when bound to agonist plus LY2119620, a new M202^5.54^ peak is observed between the peaks observed for M2R bound to LY2119620 alone and agonist alone, suggesting the formation of a distinct intermediate state. Additionally, a smaller peak is observed at a similar position to the peak in the apo-state. These data suggest that for agonist plus LY2119620, the receptor undergoes slow conformational exchange between an inactive state and an intermediate state, where the intermediate state is different from that of M2R bound to agonist alone.

**Fig. 5.**
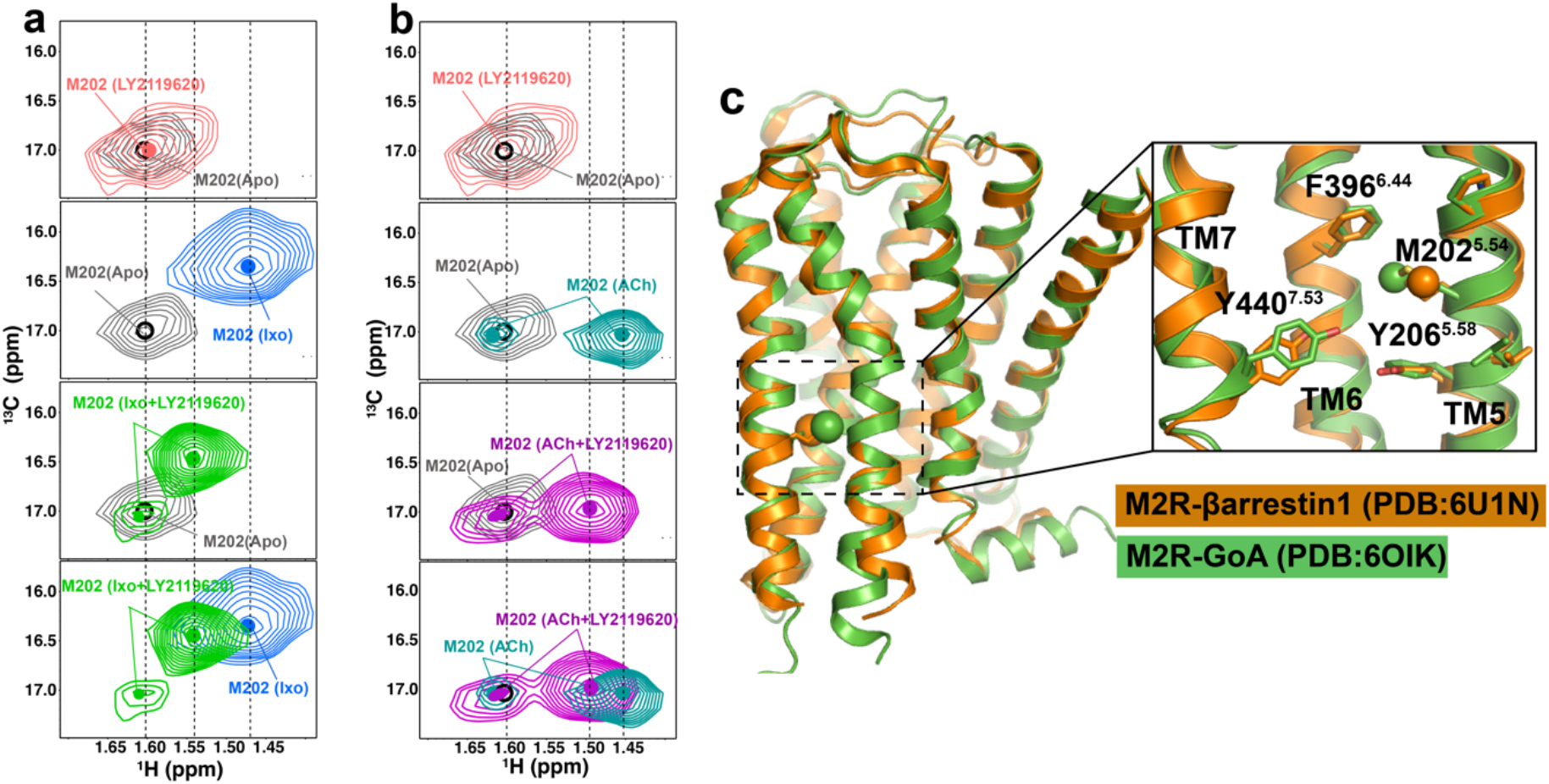
Effects of LY2119620 on the ^1^H-^13^C HSQC spectra of M202^5.54^. (a and b) The HSQC spectra of M202^5.54^ in agonist (Ixo or ACh)-bound or agonist (Ixo or ACh) + LY2119620 bound state in comparison with the apostate or LY2119620-bound state. (c) Superimposition of M2R active structures bound to GoA (green) and β-arrestin (orange). M202^5.54^ and the surrounding conserved aromatic residues Y440^7.53^, Y206^5.58^ and F396^6.44^ are shown as sticks.

It should be noted that LY2119620 appears to shift the agonist bound intermediate state toward a more inactive state when considering the chemical shifts of M202^5.54^. LY2119620 has a have a smaller effect on the ACh-bound spectrum than Ixo-bound spectrum, as a result, the ACh and Ixo spectra are more similar in the presence of the PAM (Fig. 5**a-b**). This could explain the reduced G-protein activation efficacy at high concentration of LY2119620 (Fig. 1 **a-b** **and e-f**). Our previous data indicate that the magnitude of chemical shift of M202^5.54^ correlates with agonist signaling efficacy in the absence of the active-state stabilizing nanobody Nb9-8^18^.

In contrast to the effect of LY2119620 on the Emax for G protein singlaing, the efficacy of β-arrestin recruitment is increased at high concentration of LY2119620. Thus, it is possible that the spectral changes observed for M202^5.54^ in M2R bound to LY2119620 and agonist may reflect distinct structural changes involving TM5, TM6 and TM7 that result in enhanced interactions with a GPCR kinase (GRK) and/or β-arrestin. Indeed, recent cryo-EM structures of M2R bound to GoA and β-arrestin1 show different local conformations surrounding M202^5.54^, especially the Y440^7.53^(Fig. **5c**)^25,26^. Thus, these distinct conformational states in the intracellular domain stabilized by LY2119620 binding may account for the different signaling properties, including the reduction in maximal G protein signaling and the enhanced β-arrestin recruitment efficacy (Fig. 1). Of interest, recent molecular dynamics simulations studies of the angiotensin II type 1 receptor (AT1R) show that arrestin biased ligands favor an alternative conformation in the intracellular domain of the AT1R which is distinct 27 from the canonical active conformation^27^.

## Discussion

Due to the advances in structural biology of GPCRs, crystal structures of M1-M5 subtypes are all available, including inactive, active and PAM-bound structures^15,28–30^. However, the structural basis of allosteric modulation of mAChRs still remains poorly understood due to the lack of structures bound to other allosteric modulators, and the lack of structures of agonist-bound GPCRs without stabilizing mutations, nanobodies or G proteins. Previous biophysical studies have shown that for many GPCRs, agonist alone does not stabilize a fully active state, but instead increase the probability that the receptor will populate an active state or intermediate state. Therefore, alternative approaches are required to provide additional insights into the modulation of receptor conformational states by these allosteric ligands. NMR spectroscopy has recently provided unprecedented insights into the dynamic properties of several GPCRs in response to a range of orthsoteric ligands^31–34^, while only few studies have applied NMR to examine GPCR allostery^35^. In the present study, we investigated the conformational changes surrounding five ^13^CH_3_-methionines of M2R in response to LY2119620, utilizing a previous established NMR system^18^.

Our results show that LY2119620 can dramatically affect the structure and dynamics of M2R bound to orthosteric agonists. Except for the distinct side chain of W422^7.35^ in the ECV, crystal structures of the active M2R with or without LY2119620 are remarkably similar (Fig. S3). This suggests that the PAM may preferentially bind to and stabilize the closed (active) conformation of the ECV ^7,36^. However, our NMR data show that instead of simply stabilizing the receptor in a more active state, LY2119620 can alter the conformational states of M2R along the activation process (Fig. 3, 4 and 5). Of particular interest is the reprogrammed chemical shift fingerprints of M77^2.58^ and M406^6.54^ in the presence of LY2119620 (Fig. 3), which provide direct structure evidence to show that the PAM can reshape the conformation and dynamics of the extracellular region. This may account for its partial agonist activity as well as its cooperativity with orthosteric ligands. As LY2119620 is far away from M77^2.58^ (over 10 Å) and around 8 Å away from M406^6.54^, the peaks from these ^13^CH_3_-ε-M should not be directly affected by the ligand chemistry (Fig. 2**a**). Importantly, M77^2.58^ and M406^6.54^ are located at the opposite sides of the aromatic network (Fig. 2**a**), and this region was characterized as an allosteric network that is key to the cooperativity between allosteric and orthosteric ligands by extensive mutagenesis studies^15^.

The fact that LY2119620 alone can stabilize a distinct spectrum of M2R in both the extracellular and intracellular domains may explain its direct allosteric agonism (Fig. 1). The allosteric effects of the PAM on the intracellular side of M2R are more pronounced when an orthosteric agonist is bound (Fig. 3, 4 and 5), suggesting that the allosteric and orthosteric ligands act cooperatively to stabilize distinct conformational states of the G-protein coupling domain. These distinct receptor intracellular conformational states stabilized by LY2119620 were not revealed in the previous crystal structures^7^.

In conclusion, our studies complement previous crystallography studies^7^ by providing additional structural and dynamic insights into the modulation of M2R by the PAM LY2119620. Our results suggest that LY2119620 may function by stabilizing distinct sets of conformations not observed in the presence of orthosteric agonists alone, which may account for the different signaling behaviors of the M2R when bound to LY2119620. Structural studies of mAChRs have explained why obtaining subtype selective orthosteric ligands is extremely difficult while also suggested that allosteric modulators are more promising for achieving selectivity^15,30^. A better understanding of the receptor conformational states and dynamics regulated by allosteric modulators may facilitate the development of better allosteric therapeutics targeting this class of important cell surface receptors.

## Methods

### Protein sample preparation

The ^13^CH_3_-ε-methionine labeled M2R was expressed, labeled and purified as previously described^18^. In Brief, the S19 cells were grown in methionine-deficient medium and infected at a density of 4×10^6^ ml^−1^ with M2R baculovirus in the presence of 10 uM atropine. After adding ^13^CH_3_-ε-methionine at a concentration of 250 mg L, the S19 cells were incubated for 48 hours at 27 °C. Receptor was purified by Ni-NTA chromatography, Flag affinity chromatography and size exclusion chromatography and finally exchanged to a D2O-based buffer containing 20 mM HEPEs pH 7.5, 100 mM NaCl, 0.01% LMNG, 0.003% CHS. The receptor was finally concentrated to around 80 μM for NMR experiments.

### NMR spectroscopy

The apo-state, ACh-bound, and Ixo-bound were repeated by following previously described methods^18^. For LY2119620-bound spectra, LY2119620 was added into the apo-state receptor at a concentration of 1 mM. For LY2119620 + (ACh or Ixo)-bound spectra, LY2119620 was added into the agonist-bound sample. All NMR samples were loaded into the Shigemi microtubes for data collection at 25 °C on a Bruker Avance 800-MHz spectrometer equipped with a triple-resonance cryogenic probe. The ^1^H-^13^C heter-onuclear single-quantum coherence (HSQC) spectra were recorded with spectral widths of 12820.5 Hz in 13 the 1H-dimension (w1) and 16077.2 Hz in the ^13^C-dimension (w2) centered at 45 ppm in ^13^C-dimension. For all spectra, 512 x 128 complex points were recorded and a relaxation delay of 2 s were inserted to allow spin to relax back to equilibrium. 80 scans gave rise to an acquisition time around 12 h for each spectrum. All NMR spectra were processed using the software package NMRPipe/NMRDraw and analyzed using the program NMRViewJ. The intensity for all spectra were normalized using a natural abundance peak from a highly flexible methyl group in M2R at around 1.20 ppm in the ^1^H dimension and 18.00 ppm in the ^13^C dimension as previously described^18^.

### G-protein IP-1 assay and β-arrestin recruitment assay

Measurement of G-protein mediated activation of M2R was performed applying the IP-One HTRF® accumulation assay (Cisbio, Codolet, France) in analogy to previously described protocols^18,37^. In brief, HEK 293T cells were co-transfected with the cDNA coding for M2R and the hybrid G-proteion Ga_qi_ (Ga_q_ protein with the last five amino acids at the C-terminus replaced by the corresponding sequence of Gα_i_; gift from The J. David Gladstone Institutes, San Francisco, CA) and transferred into 384 well micro plates. On the day of the experiment, cells were preincubated with the allosteric modulator (fixed concentration in the range of 0.1 to 100 μM) for 30 min before adding the agonist (final concentration for Ixo: 0.01 pM to 1 μM, for Ach: 10 pM to 300 μM) and incubating for 90 min. Accumulation of second messenger was stopped by addition of the detection reagents (IP1-d2 conjugate and Anti-IP1cryptate TB conjugate). After further 60 min time resolved fluorescence resonance energy transfer was measured using the Clariostar plate reader (BMG, Ortenberg, Germany). The obtained FRET-signals were normalized to the maximum effect of reference ACh (100%) and vehicle (0%). Normalized concentration-response curves from four to eight experiments each done in duplicates were analyzed using the algorithms for four parameter non-linear regression implemented in PRISM 8.0 (GraphPad Software, USA) to derive EC50 and Emax values.

Determination of β-arrestin-2 recruitment was performed applying the PathHunter assay (DiscoverX, Birmingham, U.K.) which is based on fragment complementation of β-galactosidase as described^18,37^. In detail, HEK293T cells were transfected with the flag-tagged M2 receptor carrying the PKA fragment for enzyme complementation and transferred into 384 well micro plates. At the day of measurement allosteric modulator at a distinct concentration was added to the cells and preincubated for 30 min followed by addition of the agonist (for Ixo: 0.1 pM to 1 μM; for Ach: 1nM to 300 μM) and incubation for further 90 min. Determination of chemoluminescence was done with a Clariostar plate reader (BMG, Ortenberg, Germany). Data analysis was performed by normalizing the raw data relative to basal activity (0%) and the maximum effect of the reference agonist ACh (100%). Normalized curves from three to five individual experiments each done in duplicate were analyzed by non-linear regression applying the algorithms in Prism 8.0 (GraphPad, San Diego, CA) to get doseresponse curves representing average EC50 and Emax value.

Significance of Emax for the agonists Ach or Ixo alone with agonist in the presence of a distionct concentration of LY2119620 in both signaling pathways was analyzed by non paired comparison by Oneway ANOVA applying Dunnett’s multiple comparisons test in Prism 8. The threshold for significance was set as 95% confidence interval and displayed as p-value (p<0.05).

## Acknowledgement

This work was supported by the Beijing Advanced Innovation Center for Structural Biology, School of Medicine, Tsinghua University and by the German Research Foundation (GRK1910 to P.G.). All NMR experiments were performed at the Beijing NMR Center and the NMR facility of the National Center for Protein Sciences at Peking University, supported by the Grant 2016YFA0501201 from the National Key R&D Program of China to C. J.. We thank Dr. Xiangyu Liu and Dr. Jie Heng for help with critical reading of the manuscript. Brian Kobilka is a Chan Zuckerberg Biohub Investigator.

## Author contribution

J.X. designed NMR experiments, performed protein purifications, processed and analyzed NMR data. H.H. performed the G protein IP-1 assay and b-arrestin recruitment assay. Y.H. and X.N. conducted the NMR experiments and assisted with NMR data processing and analysis. P.G. supervised G protein IP-1 and b-arrestin recruitment assays. C.J. supervised all NMR experiments and data processing. B.K.K. provided overall project supervision. J.X. and B.K.K. wrote the manuscript with contributions from all authors.

## Supplementary figures

**Fig. S1.**
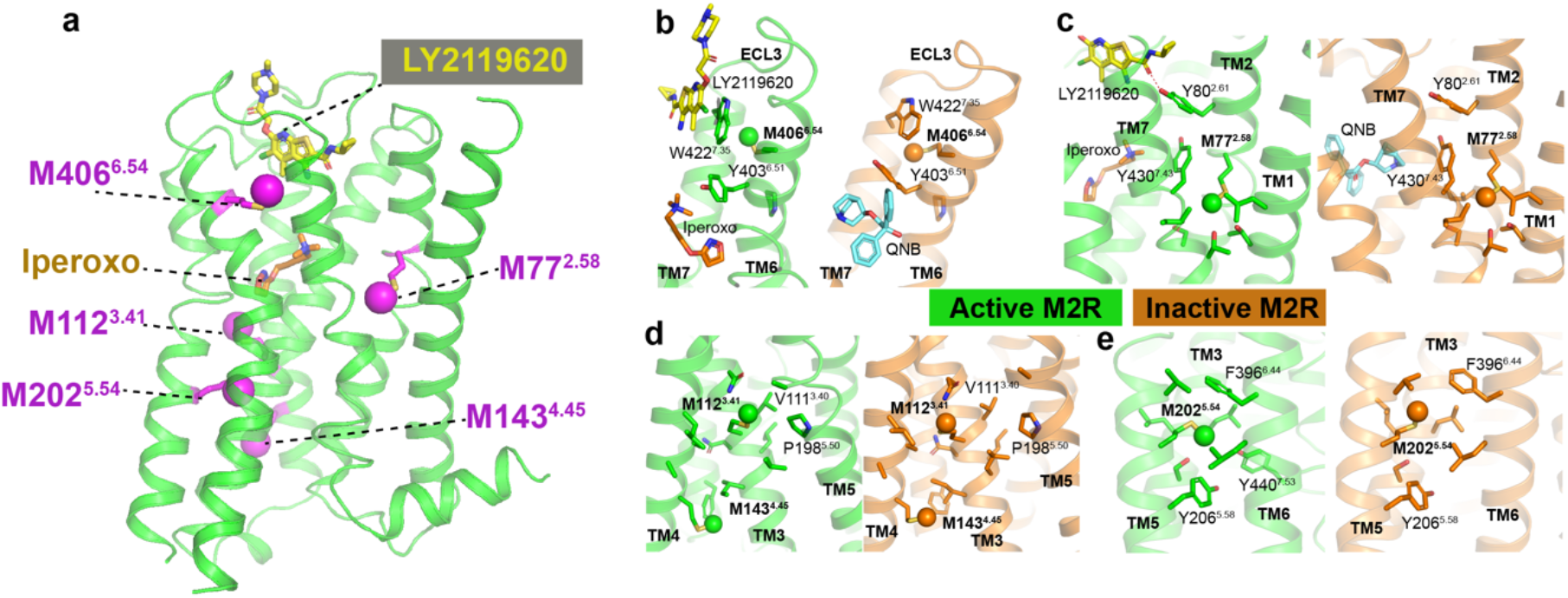
Distribution of the five ^13^CH_3_-ε-methionines in the M2R. Methionines are shown as sticks in magenta, and the ^13^C atoms were shown in sphere.

**Fig. S2.**
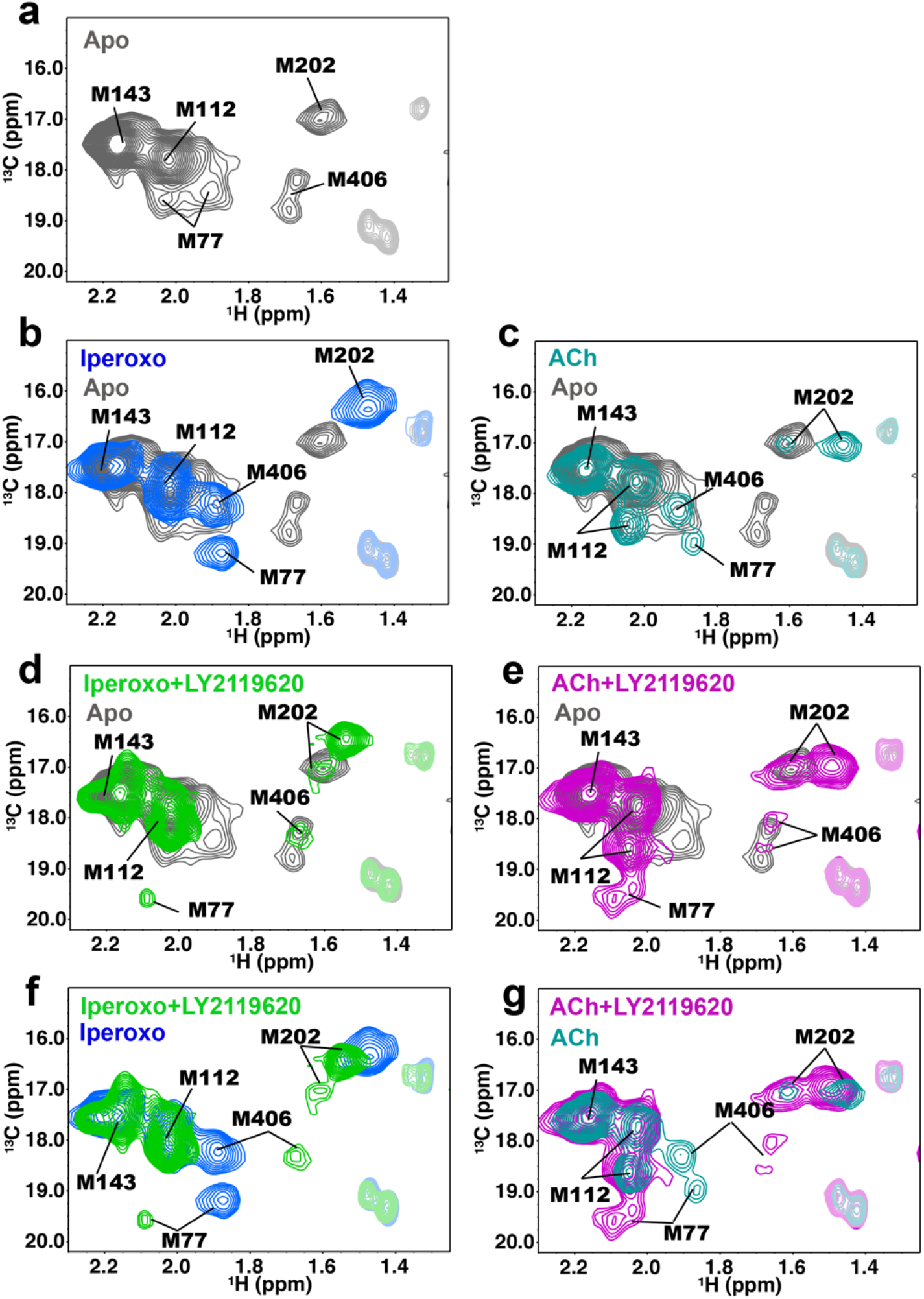
^1^H-^13^C HSQC spectra of M2R bound to different ligands or ligand combinations. (a) Spectra of apo-state M2R. (b-c) Spectra of M2R bound to Iperoxo (b) and ACh (c). (d-e) Spectra of M2R bound to Iperoxo and LY2119620 (d) and ACh and LY2119620 (e). The apo-state spectrum is set as reference and shown in gray. (f-g) Comparison of the overall spectra of agonist (iperoxo or ACh)-bound M2R in the presence or absence of LY2119620.

**Fig. S3.**
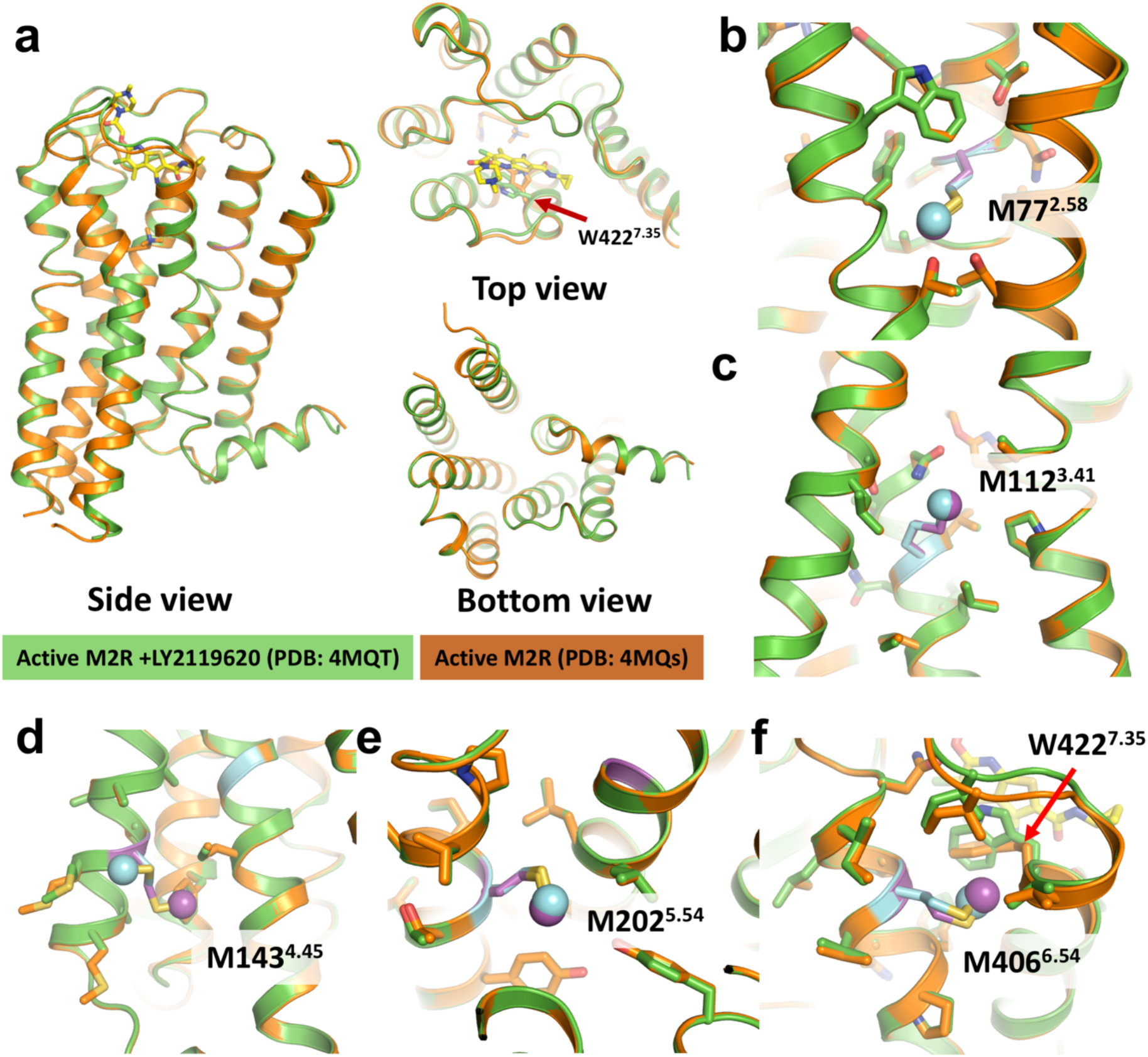
Superimposition of M2R active structures with and without LY2119620 bound reveal few difference between these two structures. (a) Comparison of the overall structures. Within the extracellular vestibule, Trp 422 ^7.35^ undergoes a change of rotamer (top view). (b-f) Comparison of the local chemical environments surrounding each ^13^CH3-ε-methionine. Residues within 4 Å of each methionine are shown in sticks. ^13^CH3-ε-methionines in active and active plus PAM structures are shown in magenta and cyan, respectively. The ^13^C atoms are shown as spheres.

**Table S1:**
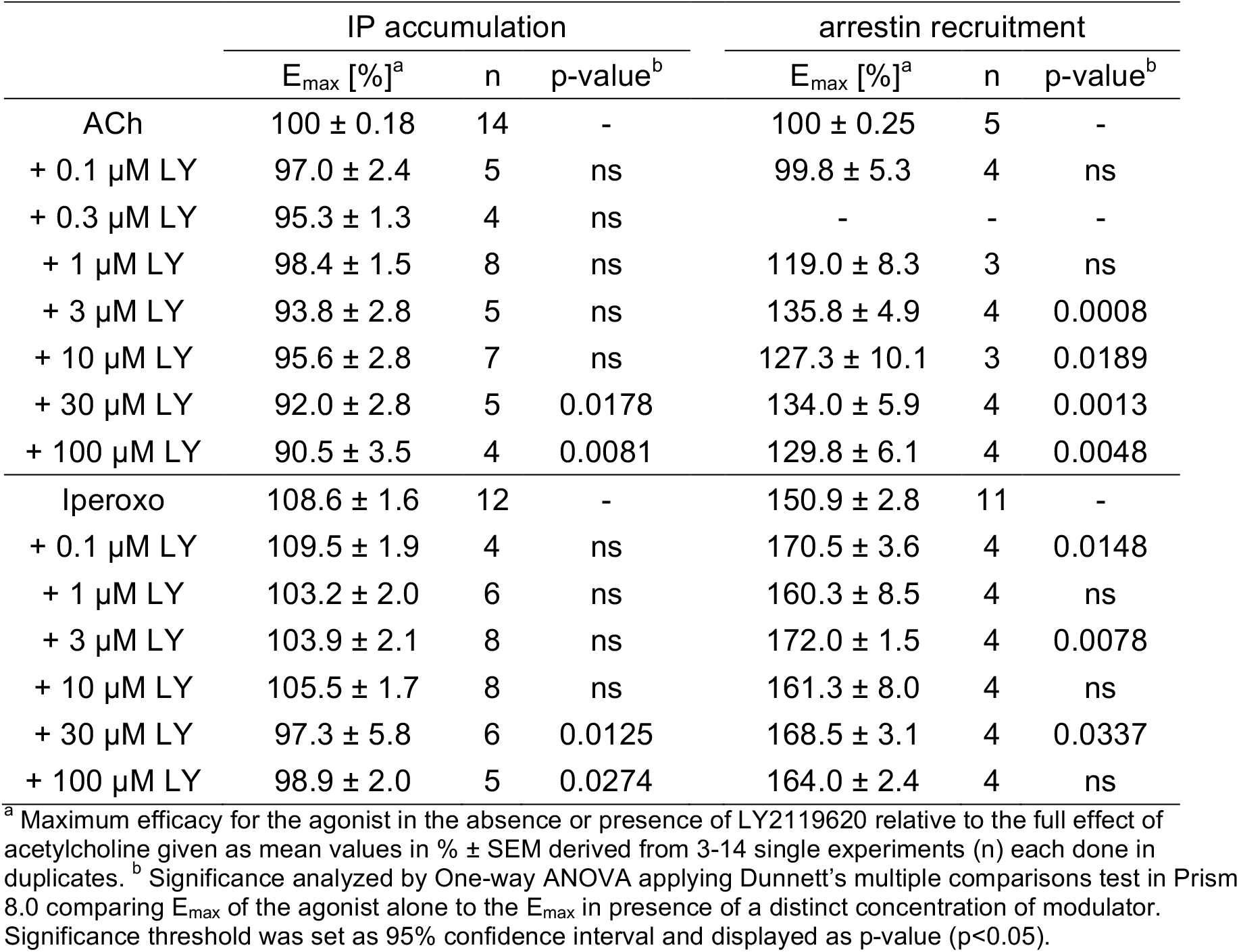
Maximum efficacy of ACh and Iperoxo in the absence and presence of the modulator LY2119620 at M2R for G-protein activation measured by a cell-based Gqi-inositol phosphate accumulation assay and ß-arrestin-2 recruitment determined by the PathHunter assay.

